# A Hypothesis-Based Hop Microbiology Laboratory Module Testing the Plausibility of the Mythical Origin of the India Pale Ale (IPA)

**DOI:** 10.1101/2023.07.27.550083

**Authors:** Julio Molina Pineda, Amanda N. Scholes, Jeffrey A. Lewis

## Abstract

As one of the most famous fermented drinks in the world, beer is an especially relatable topic for microbiology courses. Here, we describe a short and easily adaptable module based on the antibacterial properties of hops used in brewing. By the 15^th^ century, beer recipes included hops (the flower of the *Humulus lupulus* plant) as a bittering agent and antimicrobial. By the 19^th^ century, the highly-hopped Indian Pale Ale (IPA) became popular, and a modern myth has emerged that IPAs were invented to survive long ocean voyages such as from Britain to India. With that myth in mind, we designed a hypothesis-driven microbiology lab module that tests the plausibility of this brewing myth— namely that highly-hopped beers possess enough antibacterial activity to prevent spoilage, while lowly-hopped beers do not. The overall design of the module is to test the antimicrobial properties of hops using petri plates containing varying concentrations of hop extract. The module includes hypothesis generation and testing related to bacterial physiology and morphology (hops are not equally effective against gram-positive and gram-negative bacteria), and to mechanisms of antimicrobial resistance (as beer spoilage bacteria have repeatedly evolved hop resistance). Pre and post assessment showed that students made significant gains in the learning objectives for the module, which encourages critical thinking and hypothesis testing by linking microbial physiology and antimicrobial resistance to an important and topical real-world application.

## Introduction

Beer is rich in nutrients, yet spoilage is rare. In part, this is due to the antibacterial effects of ethanol, but bittering agents play a key role. Up until the Middle Ages, beer was bittered with gruit, a mixture of herbs mainly consisting of sweet gale and bog myrtle (1). By the 14^th^ century, gruit was ultimately supplanted by hops, the flowers from the plant *Humulus lupulus*, as hops were cheaper and more consistent as a beer preservative (1, 2). In the 19^th^ century, highly-hopped beers such as the Indian Pale Ale (IPA) came into vogue, with a matter of historical dispute arising over their origin (3, 4). A common (though likely untrue) myth is that IPAs were popularized due to their ability to survive long ocean voyages (such as Britain to India, hence the name). With that myth in mind, we designed a hypothesis-driven module for a college microbiology lab course that tests the plausibility of the myth—namely that highly-hopped beers possess enough antibacterial activity to prevent spoilage, while lowly-hopped beers do not.

The overall design of the module is to test the antimicrobial properties of hops in a representative gram-positive and gram-negative bacterium on petri plates containing varying concentrations of hop extract. Hop iso-alpha acids are responsible for both bitter flavor and the antibacterial activity of hops, and are thought to act as ionophores that disrupt proton gradients (5, 6). Gram-negative bacteria are naturally hop resistant, likely due to the impermeability of the outer membrane to iso-alpha acids (7, 8). Thus, an added aspect of this module is discussion of bacterial cell wall morphology and generating hypotheses for whether hop antimicrobial activity would be more effective against gram-positive or gram-negative bacteria. Interestingly, despite the fact that gram-negative bacteria are naturally resistant to hops, the vast majority of spoilage bacteria are gram-positive bacteria (7). The evolution of hop resistance in gram-positive bacteria is one major reason for this disparity (7, 8), and the final part of the module is a group discussion to generate plausible hypotheses for mechanisms of evolved hop resistance. Overall, this relatable module engages students in fundamental concepts in bacterial physiology, antibacterial activity, and the evolution of antibacterial resistance.

### Learning Objectives

This 2-part module (1 week between lab periods) was originally piloted as a standalone activity while waiting for beer to ferment for an upper-level lab course on the microbiology of brewing (9), and was refined to be adaptable to any college-level microbiology lab course.

Following completion of the module, students should be able to:

LO1. Describe the history and importance of hops in brewing, including the active chemicals responsible for bitterness and antimicrobial activity.

LO2. Ascertain the plausibility of the ‘myth’ that highly-hopped India Pale Ales (IPAs) were developed specifically to prevent spoilage during long ocean voyages.

LO3. Describe the differences in cell wall morphology between gram-positive and gram-negative bacteria, and the implications for hop sensitivity.

LO4. Understand the likely cellular mechanism of iso-alpha acids toxicity, and predict the type of genes that are targets for hop-resistance mutations in beer-spoilage bacteria.

## Materials and Procedure

### Safety Issues

Biosafety level 1 (BSL-1) category bacterial strains used were non-pathogenic *Escherichia coli* K12 strain MG1655 and a food-grade strain of *Lactobacillus buchneri*. During all lab activities, students should wear personal protective equipment (safety glasses, lab coats, gloves). Concentrated iso-alpha acid extract is a moderate hazard (skin and eye irritant) and may cause allergic skin reactions.

### Preparations Before the Module: Media and Bacterial Culturing

Lysogeny Broth (LB) agar (for *E coli*) and Lactobacilli MRS Broth (MRS) agar (for *L. buchneri*) are prepared using 4 different concentrations of iso-alpha acids: 0 International Bittering Units (IBU) (control), 10 IBU (lager level of hops), 50 IBU (IPA level of hops), and 100 IBU (double or imperial IPA level of hops). We empirically determined via octanol extraction and UV-Vis spectroscopy (10, 11) that 1 IBU per L media corresponded to 26 μl of 30% (w/w) iso-alpha acid extract. Iso-alpha acid extract is added to media prior to autoclaving (20 min at 120°C). Plates should be protected from light (iso-alpha acids are UV sensitive) and stored at 4°C for up to 1 week before use.

One-week before the experiment, streak out bacteria from frozen stocks (20% glycerol) onto solid media. *E. coli* colonies arise after 1 day at 37°C and can be stored at 4°C until propagating overnight to saturation in liquid LB at 37°C with orbital shaking (270 rpm). *L. buchneri* colonies arise after 72 hours at 37°C, and are propagated to saturation in liquid MRS at 30°C without shaking for 48 hours (12). On the day of the experiment, instructors prepare 1-ml working stocks for each student. For *E. coli*, saturated cells were diluted to an OD_600_ of 0.5 and then were further diluted 10^-4^ (∼ 4 x 10^5^ cells/ml). For *L. buchneri*, cells were diluted to an OD_600_ of 0.3.

### Procedures during the Module: Part 1

The first part of the module consists of a short lecture on the history of hops, the mythical origins of IPAs, bacterial cell wall physiology, which is then followed by a group discussion on hypotheses about whether gram-positive or gram-negative bacteria are more likely to be affected by hops. Following the short lab lecture, students then prepare serial dilutions of bacteria to spot onto the plates with different levels of iso-alpha acids. Each student is provided with their 1-ml working stocks of bacteria, 1 ml each of sterile LB and MRS broth, and a sterile 96-well plate. Students prepare serial dilutions in the 96-wll plate by transferring and mixing 10 μl of culture to 90 μl of sterile media, followed by spotting 4 μl (*L. buchneri*) or 20 μl (*E. coli*) of the final (10^-4^) dilution onto the appropriate plates. *E. coli* plates are incubated at 37°C for 24 hours, while *L. buchneri* plates are incubated at 37°C for 72 hours. Once colonies have grown on the control plates, the plates can be stored at 4°C until the second part of the module.

### Procedures during the Module: Part 2

The second part of the module begins with students interpreting their results in light of their original hypotheses. Figure 1 depicts representative results, with the gram-negative *E. coli* being fully resistant to iso-alpha acids and gram-positive *L. buchneri* showing growth only at low (10 IBU) levels of iso-alpha acids. These results illustrate that hops specifically inhibit gram-positive bacteria, while also providing some credence to the notion that IPA-levels of hops could reduce beer spoilage during long ocean voyages. Finally, it is pointed out that gram-positive bacteria can evolve hop resistance, which is still a problem for industrial brewers, and students discuss as a group plausible molecular mechanisms for how hop resistance may arise.

**Figure 1.**
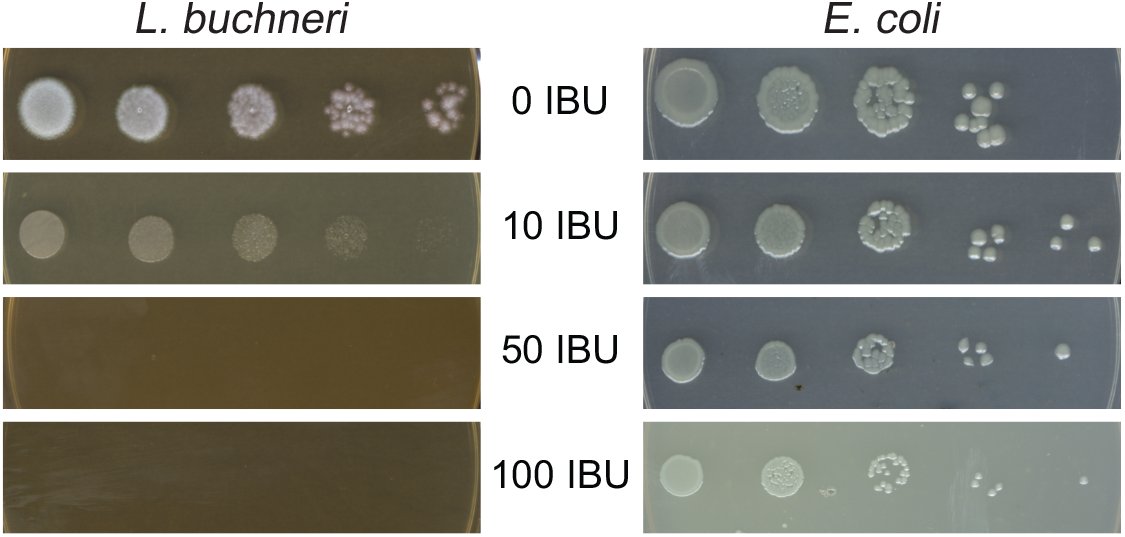
Representative results of serial dilutions of *L. buchneri* and *E. coli* on plates containing the indicated levels of iso-alpha acids.

### Assessment

Student learning was assessed via graded lab notebooks, and pre and post exams during the Fall of 2022. The average pre-test score was 20% correct, which rose to 60% following completion of the module (Figure 2, n = 10). These gains were both statistically significant (*P* = 0.002, Wilcoxon signed-rank test) and of large effect (Cliff’s Delta = 0.88). Notably, students performed better upon completing the module on questions relating to the biological reasons for differences in hop sensitivity and hypothesizing potential mechanisms for hop resistance arising (Figure 3), highlighting that this module helps with teaching advanced concepts in microbial physiology and genetics.

**Figure 2.**
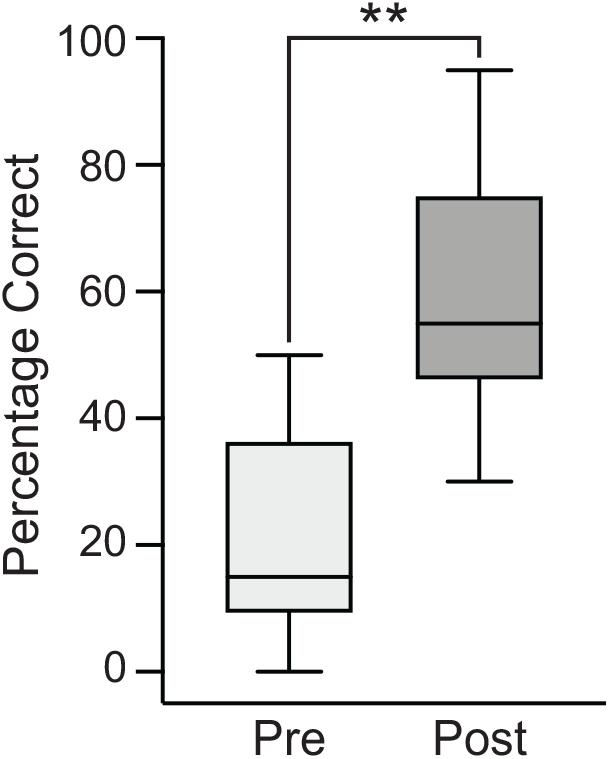
Pre- and post-exams show significant gains in student learning. The boxplot depicts the median and interquartile range of overall exam scores, while the whiskers denote the range. **, *P* = 0.002, Wilcoxon signed-rank test.

**Figure 3.**
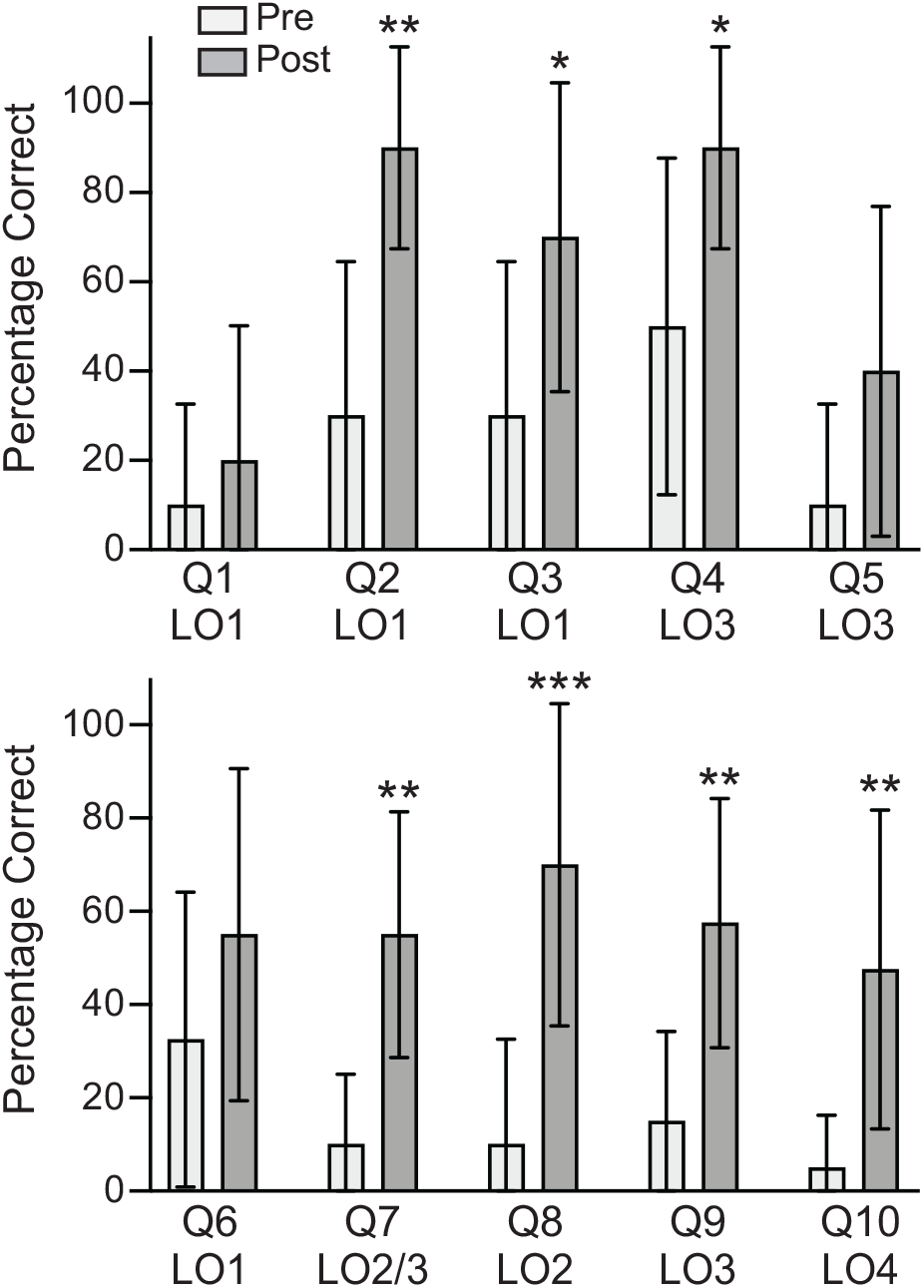
Bar graph depicting the mean and 95% confidence intervals for individual item responses for pre- and post-exams. Exam questions can be found in Appendix 2. LO denotes the learning objectives; *, *P* < 0.05; **, *P* < 0.01; ***, *P* < 0.001, two-way ANOVA, Fisher’s LSD test.

## Discussion and Conclusions

We created a microbiology lab module that can be easily adapted and integrated into a college-level microbiology or molecular biology lab course. Our pre- and post-test assessment provides evidence that students made significant gains in our learning objectives. Overall, this module encourages critical thinking and hypothesis testing by linking microbial physiology and antimicrobial resistance to an important and topical real-world application.

## Acknowledgments

Funding for this project was supported in part by National Science Foundation grants IOS-1656602 and MCB-1941824. We thank Chenguang Fan for providing us with *E. coli* K12 MG1655 and Christian Tipsmark for the use of equipment. The authors declare that there are no conflicts of interest.

