## Appendices for "A Hypothesis-Based Hop Microbiology Laboratory Module Testing the Plausibility of the Mythical Origin of the India Pale Ale (IPA)"

### Supplementary Material

Appendix 1: Supplies and Consumables

Appendix 2: Pre- and Post- Assessment Quiz Questions

Appendix 3: Media Recipes

Appendix 4: Lab Lectures

Appendix 5: Student Instructions Handout

**Appendix 1. Supplies and Consumables**

| Item | Company | Catalog Number | Quantity (for 12 students) |
|---|---|---|---|
| Yeast extract | VWR | 90000-726 | 1 bottle (500 g) |
| Tryptone | VWR | 97063-390 | 1 bottle (500 g) |
| NaCl | VWR | BDH9286-500G | 1 bottle (500 g) |
| Lactobacilli MRS Broth | Criterion | C5931 | 1 bottle (500g) |
| Agar | Fisher | BP1423-500 | 1 bottle (500 g) |
| Iso-alpha-acids 30% w/w (IsoHop) | Willamette Valley Hops | NA | 1 bottle (1 kg) |
| *Lactobacillus buchneri*, food grade | Wyeast | 5335 | 1 packet |
| Flasks (for pouring media) | NA | NA | 8 |
| Small petri plates | VWR | 25384-342 | 100 plates |
| P10 tips | VWR | 89168-748 | 1 case |
| P200 tips | VWR | 89140-900 | 1 case |
| Gloves (various sizes) | Santa Cruz Biotech | sc-359596 | 1 box per size |
| Ethanol burners/Bunsen burner | NA | NA | 12 |
| 95% Ethanol (for burners) | VWR | BDH1158-4LP | 1 bottle (4L) |
| P10 pipette | MidSci | A-10 | 12 |
| P100 pipette | MidSci | A-100 | 12 |
| 96-well plate | VWR | 76210-520 | 12 |
| 1.7 ml microcentrifuge tubes | VWR | 89126-724 | 1 box of 500 |
| Tabletop incubators | NA | NA | 1 at 30°C and 1 at 37°C |
| Shaking incubator | NA | NA | 1 at 37°C |
| Autoclave | NA | NA | 1 |

**Appendix 2. Pre- and Post- Assessment Quiz Questions**

**Unique Identifier (birth month, mother's first initial, birth day):**

*Multiple Choice:*

**1.** All of the following are true regarding hops ***except***…

A. Hops are generally added during the boiling step of brewing.
B. Hops refers to the flower of the *Humulus lupulus* plant.
C. Hops have broad antimicrobial activities against both eukaryotic and prokaryotic spoilage microbes.
D. Hops largely replaced an herb mix called "gruit" in brewing by the end of the Middle Ages.
E. All of the above are true.

**2.** The main type of chemical compound in hops responsible for beer bitterness is…

A. flavonoids
B. polyphenols
C. terpenes
D. iso-alpha acids
E. thioesters

**3.** The main type of chemical compound in hops responsible for antimicrobial activity is…

A. flavonoids
B. polyphenols
C. terpenes
D. iso-alpha acids
E. thioesters

**4.** Which of the following is a ***major difference*** between gram-positive and gram-negative bacteria?

A. Gram-positive bacteria possess a thick cell wall composed predominantly of peptidoglycan.
B. Gram-positive bacteria possess an outer membrane that contains lipopolysaccharide (LPS).
C. Gram-positive bacteria are rarely the causative agents of beer spoilage.
D. Gram-negative bacteria possess a lightly staining nucleus.
E. Gram staining of gram-negative bacteria yields cells with a deep violet color.

**5.** What is thought to be the main molecular mechanism of hop compounds' antimicrobial activity?

A. They act as inhibitors of peptidoglycan biosynthesis.
B. They act as inhibitors of lipopolysaccharide (LPS) biosynthesis.
C. They act as inhibitors of RNA polymerase.
D. They act as nucleases that degrade cellular nucleic acids.
E. They act as ionophores that disrupt proton gradients.

*Short Answer Questions:*

**6.** Hops can be optionally added during the secondary fermentation in what is called "dry hopping." Would this type of hop addition impart any additional bitterness? Why or why not?

**7.** Sour beers are frequently made by fermenting with acid-producing *Lactobacillus sp.* in addition to yeast. Approximately what IBU level would you predict those beers to be, and why?

**8.** One ‘myth’ about the origin of India Pale Ales (IPAs) is that they were developed to avoid spoilage during the long voyage from England to India. Based on the expected antimicrobial activity of IPA-level hop concentrations, is this myth incorrect or “busted?” Why or why not?

**9.** Are hops more effective against gram-positive or gram-negative bacteria, and what is responsible for the difference?

**10.** Hop resistant mutants arise readily in nature, which is one reason that brewers still worry about spoilage. Explain one possible mechanism for evolving hop resistance. In other words, what types of genes might be affected by resistance mutations, and how do those effects confer hop resistance?

*Multiple Choice:*

**1.** All of the following are true regarding hops ***except***…

A. Hops are generally added during the boiling step of brewing.
B. Hops refers to the flower of the *Humulus lupulus* plant.
C. Hops have broad antimicrobial activities against both eukaryotic and prokaryotic spoilage microbes.
D. Hops largely replaced an herb mix called "gruit" in brewing by the end of the Middle Ages.
E. All of the above are true.

**2.** The main type of chemical compound in hops responsible for beer bitterness is…

A. flavonoids
B. polyphenols
C. terpenes
D. iso-alpha acids
E. thioesters

**3.** The main type of chemical compound in hops responsible for antimicrobial activity is…

A. flavonoids
B. polyphenols
C. terpenes
D. iso-alpha acids
E. thioesters

**4.** Which of the following is a ***major difference*** between gram-positive and gram-negative bacteria?

A. Gram-positive bacteria possess a thick cell wall composed predominantly of peptidoglycan.
B. Gram-positive bacteria possess an outer membrane that contains lipopolysaccharide (LPS).
C. Gram-positive bacteria are rarely the causative agents of beer spoilage.
D. Gram-negative bacteria possess a lightly staining nucleus.
E. Gram staining of gram-negative bacteria yields cells with a deep violet color.

**5.** What is thought to be the main molecular mechanism of hop compounds' antimicrobial activity?

A. They act as inhibitors of peptidoglycan biosynthesis.
B. They act as inhibitors of lipopolysaccharide (LPS) biosynthesis.
C. They act as inhibitors of RNA polymerase.
D. They act as nucleases that degrade cellular nucleic acids.
E. They act as ionophores that disrupt proton gradients.

*Short Answer Questions:*

**6.** Hops can be optionally added during the secondary fermentation in what is called "dry hopping." Would this type of hop addition impart any additional bitterness? Why or why not?

No, because the lack of boiling would not isomerize the alpha acids.

**7.** Sour beers are frequently made by fermenting with acid-producing *Lactobacillus sp.* in addition to yeast. Approximately what IBU level would you predict those beers to be, and why?

10 IBU or less because *Lactobacillus sp.* are likely hop sensitive.

OR

IBU >10 because those particular *Lactobacillus sp.* have evolved hop resistance.

**8.** One 'myth' about the origin of India Pale Ales (IPAs) is that they were developed to avoid spoilage during the long voyage from England to India. Based on the expected antimicrobial activity of IPA-level hop concentrations, is this myth incorrect or "busted?" Why or why not?

No, the myth is plausible because IPA-levels of hops strongly inhibited one of the species tested.

**9.** Are hops more effective against gram-positive or gram-negative bacteria, and what is responsible for the difference?

Gram-positive, likely because of differences in the cell membrane architecture.

**10.** Hop resistant mutants arise readily in nature, which is one reason that brewers still worry about spoilage. Explain one possible mechanism for evolving hop resistance. In other words, what types of genes might be affected by resistance mutations, and how do those effects confer hop resistance?

Known mutations that confer hop resistance are those that increase expression of efflux pumps, increase expression of plasma membrane ATPase (help restore proton motive force), or increase expression of $Mn^{+}$ importers. Other plausible mechanisms would be mutations that allow an enzyme to modify iso-alpha-acids so that they no longer function as ionophores, or mutations that cause a protein to bind and thus "tie up" iso-alpha-acids, or mutations that affect cell membrane composition so that iso-alpha-acids no longer easily enter the cell.

**Appendix 3. Media Recipes**

**Caution: concentrated hop extract is a moderate hazard (skin and eye irritant) and may cause allergic skin reactions.**

**Note: the amounts of hop extract were empirically determined and may need to be adjusted. See Figure 1 for expected results, and optimize if necessary.**

**Lysogeny Broth (LB) for *E. coli*:**

1. Mix the following in 500 ml of distilled water:

    5g Yeast Extract

    10g Tryptone

    5g NaCl

2. Stir until fully dissolved, then bring volume to 1 L.

3. For plates only:

    3a. Add corresponding amount of 30% iso-alpha-acid extract based on 26 µl per L per IBU (e.g., 260 µl for 10 IBU plates).

    3b. Add 15g of agar, stir for 5 minutes.

4. Autoclave at 120°C for 20 min.

**De Man, Rogosa, and Sharpe Broth (MRS) for *L. buchneri*:**

1. Mix 55g of Criterion Lactobacilli MRS Broth (C5931) with 500 ml of distilled water.

2. Stir until fully dissolved, then bring volume to 1 L.

3. For plates only:

    3a. Add corresponding amount of 30% iso-alpha-acid extract based on 26 µl per L per IBU.

    3b. Add 15g of agar, stir for 5 minutes.

4. Autoclave at 120 °C for 20min.

**Appendix 4. Lab Lectures**

**Student responses to discussion prompts are included in bolded, red text.**

**Week 1 Lecture:**

**Slide 1**

(Title) Preservatives in Beer Brewing

- Grain extracts have less flavor compared to the fruit used to make wine, so ancient brewers added spices as additional flavoring agents
- Ancient fermentations relied on airborne yeast for inoculation
  - lower starting amounts of yeast increases the chance of spoilage microbes taking over
- Beer is also more perishable than wine due to much lower alcohol content
- Thus, spices used for beer flavoring were also chosen based on how well they acted as preservatives

**Slide 2**

Hop History

- Prior to ~800AD, beer was bittered with a herb and spice mixture known as gruit.
  - mainly sweet gale (*Myrica gale*) and also rosemary, ginger, spruce, juniper, and others
  - Pronounced as "groot" ('grüt)
- Gruit did have preservative properties, though inconsistent
- Hops (*Humulus lupulus*) are climbing plants native to Europe (as well as western Asia and North America).
- First documented cultivation of hops was 736 AD, and it became commercially cultivated in Germany in the 1100's.
- 1400's: hopped beer brewed in England, and hops overtakes gruit as it is cheaper and more consistent as a preservative
- 1710: English parliament banned the use of non-hop bittering agents

**Slide 3**

Hop Botany

- cones of the female *Humulus lupulus* plant
  - Image: hop cones
- Resins (alpha and beta acids) and essential oils are present in unique organs called lupulin glands
  - Image: cross section of cut hop cone showing lupulin glands

**Slide 4**

Boiling wort with hops...

- isomerizes the alpha acids in the resins (generating iso-alpha-acids)
  - e.g., humulone (most abundant alpha acid in hops)
  - Image: humulone isomerization reaction

**Slide 5**

Iso-alpha Acids

- are largely responsible for hop bitterness
  - bitterness is measured in International Bittering Units (IBU)
  - 1 IBU = 1 mg of iso-alpha-acids per liter
- are also responsible for the anti-microbial activity of hop addition

**Slide 6**

Hop Levels of Different Beer Styles

- Table: A General IBU Guide
  - Call outs of American Light Lager 8-12 IBU [10 IBU],
  - Indian Pale Ale (IPA) 60-80 IBU [50 IBU],
  - Double or Imperial IPA 80-100 IBU [100 IBU]

**Slide 7**

The (Somewhat Mythical) History of the India Pale Ale

- Sometimes attributed to the brewer George Hodgson, who made a heavily hopped aged beer that could survive the 4-6 month journey to India (late 1780's)
- Pale ales were preferred over porters in tropical climates
- By 1760's brewers were advised to add extra hops to beers being exported to warmer climates
- By 1822, Pale Ales for the India market were described as having twice the usual quantity of hops
- From 1841 onwards, IPAs became increasingly popular in Britain
- Mild US resurgence in the 1970's and then major in the 1990's, with new styles emerging in the 2000's (e.g., New England hazy IPA)

**Slide 8**

Beer Spoilage Microbes

- ~75% of beer spoilage microbes are gram-positive lactic acid bacteria (mostly *Lactobacillus* and *Pediococcus* species).
- Remainder are wild yeast (e.g., Brettanomyces—leads to "funky" flavor) or gram-negative bacteria (that mostly produce acetic acid (vinegar))

**Slide 9**

Would you expect the anti-microbial activity of hop iso-alpha acids to work against wild yeast?

- **Mixed discussion that settled on "no," because if hops inhibited wild yeast, they would likely inhibit the growth of brewing yeast as well.**

**Slide 10**

There are two major morphological classes of bacteria: gram-positive and gram-negative

- Image: differences in morphology of gram positive and gram negative cell wall, showing specifically the large peptidoglycan layer of gram-positive bacteria, and the thin peptidoglycan layer and lipopolysaccharide outer membrane of gram-negative bacteria
- Verbally point out that the nomenclature is based on the Gram stain, and why gram-positive and gram-negative bacteria are differentially stained

**Slide 11**

Image: Phylogenetic tree showing major classes of gram-positive and gram negative bacteria

- Arrows specifically call out *Escherchia* and *Lactobacillus* on nearly opposite ends of the tree

**Slide 12**

Do you expect the anti-microbial activity of hop iso-alpha acids to work equally well against Gram-positive or Gram-negative bacteria?

- **Mixed discussion with split answers: yes, both are bacteria or no, the differences in cell wall composition likely causes differences in hop effectiveness**

**Lab Activity: make serial dilutions and spot bacteria on control plus hop plates.**

**Week 2 Lecture:**

**Lab Activity: examine plates and record results. Once completed instructors proceed to the lab lecture.**

**Slide 1**

Hop Levels of Different Beer Styles

- Table (from week 1): A General IBU Guide
  - Call outs of American Light Lager 8-12 IBU [10 IBU],
  - Indian Pale Ale (IPA) 60-80 IBU [50 IBU],
  - Double or Imperial IPA 80-100 IBU [100 IBU]

**Slide 2**

*Lactobacillus buchneri* is commonly-used to make sour beers (we bought our strain from White labs). If you were using *L. brevis* to make a sour, what IBU would you target for your wort?

- **Students argued for 5-10 IBU based on the poor growth of *L. buchneri* above 10 IBU**

**Slide 3**

Image: Graph of Percent Growth of *buchneri* from White Labs, showing severe reduction of growth beyond 10 IBU

**Slide 4**

The Mythical Origin of IPAs, Small Group or Individual Discussion Question

- "Highly hopped beer was made to survive the 4–6-month journey from Europe to India."
- So, is this myth plausible, or is it "busted?"
  - **Students argued plausible because 50 IBU strongly inhibited the gram-positive bacterium (and the majority of spoilage microbes are Gram-positive)**

**Slide 5**

There are two major morphological classes of bacteria: gram-positive and gram-negative

- Image (from week 1): differences in morphology of gram positive and gram negative cell wall,
- Remind students of the two major morphological classes of bacteria and their difference in cell wall composition.

**Slide 6**

- Do hops appear more effective against Gram-positive or Gram-negative bacteria?
  - **Students all answered "gram-positive bacteria"**
- What are some limitations to this experiment?
  - **Student answers included: only tested one hop component, did not test in beer (which has ethanol), and only tested two different species (so may not be completely representative of all gram-positive and gram-negative bacteria)**

**Slide 7**

The Mechanism of Iso-alpha acid anti-Bacterial activity

- Act as "ionophores" against gram-positive bacteria
- Image: cartoon of carrier vs channel ionophores
- Image: iso-humulone's chemical structure
- Do you expect iso-humulone to act as a carrier or channel-forming ionophore?
  - **Students answered "carrier" based on it not looking like a pore/channel**

**Slide 8**

The Mechanism of Iso-alpha acid anti-Bacterial activity

- The gram-negative outer membrane is highly effective at keeping out polar compounds (like iso-alpha-acids).
- Gram-positive bacteria lack this outer membrane and are thus more susceptible to ionophores.

**Slide 9**

The Mechanism of Iso-alpha acid anti-Bacterial activity

- Iso-alpha-acids are protonated and release $H^+$ into the cytoplasm, acidifying it.
- Iso-alpha acids also bind to $Mn^+$ ions, leading to export from the cytoplasm
- Overall, disrupts proton motive force, enzyme activity, and transport (e.g., nutrients)
- Image: Left half of Figure 4 from Bokulich NA and Bamforth CW. 2013. Microbiol Mol Biol Rev. 77:157-172.

**Slide 10**

- Contamination with gram-positive lactic acid bacteria is still common.
- So, bacteria must have evolved resistance mechanisms.
- How might cells have evolved resistance?
    - **Student answers included: pump protons out the cell in response to iso-alpha acids, evolve proteins to bind up iso-alpha-acids, and toughen cell wall so that iso-alpha acids cannot enter**

**Slide 11**

Actual Resistance Mechanisms

- plasmid-encoded HorA and/or ORF5 (multi-drug transporters).
- upregulated expression of *hitA*, increasing $Mn^+$ influx
- upregulated expression of plasma membrane ATPase (increase $H^+$ efflux, help restore proton motive force)
- Image: Full Figure 4 from Bokulich NA and Bamforth CW. 2013. Microbiol Mol Biol Rev. 77:157-172.

**Appendix 5. Student Instructions Handout**

**Hop Sensitivity Assay**

The alpha acids in hop flowers are isomerized during wort boiling to form **iso-alpha-acids**. Humulone shown below is the most prevalent alpha acid in hops.

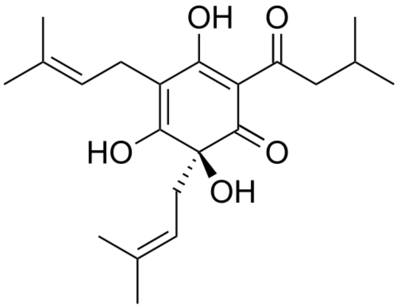


humulone

The hop iso-alpha-acids contribute bitterness to the beer, while also having **antimicrobial** effects. The amount of iso-alpha acids is generally reported on the **International Bittering Units** (IBU) scale, with 1 IBU equaling 1 mg of iso-alpha-acids per liter of beer. IBUs for common beer styles range from less than 10 (e.g., lagers, lambics) to over 100 ("imperial" IPAs and stouts, barley wines). The goal of this assay is to measure hop sensitivity of a gram-positive (*Lactobacillus buchneri*) and gram-negative (*Escherichia coli*) bacterium at low (10 IBU: light lagers), moderate (50 IBU: IPAs), and high levels (100 IBU: imperials) of hop iso-alpha acids.

**Preparation by TA:**

1. Grow cultures of *L. buchneri* (Gram-positive) and *E. coli* (Gram-negative) to saturation at 30°C and 37°C, respectively. Dilute *E. coli* to an $OD_{600}$ of 0.5 and then further dilute $10^{-4}$ (~ 4 x $10^5$ cells/ml). Dilute *L. buchneri* to an $OD_{600}$ of 0.3.

2. For *L. buchneri* prepare Lactobacilli MRS broth plates with varying amounts of iso-alpha-acid extract (0 IBU control, 10 IBU, 50 IBU, 100 IBU).

3. For *Escherichia coli* prepare LB plates with varying amounts of iso-alpha-acid extract (0 IBU control, 10 IBU, 50 IBU, 100 IBU).

**Hop Sensitivity Assay:**

1. In the provided 96-well plate, add 90 µl of LB to each of 4 consecutive wells of one row, leave the first well of the row empty.

2. Add 90 µl of MRS to each of 4 consecutive wells of a second row, leave the first well of the row empty.

3. Starting with *E. coli*, briefly vortex the culture tube and transfer 100 µl to the first well of the **LB row**. Then, transfer 10µl from this well into the first well with media on it and mix by pipetting with a P-100 pipettor. Since you have added 10 µl of cells to 90 µl of media, this is a 1:10 or $10^{-1}$ dilution. **Note: make sure that *E. coli* is added to LB and not MRS media.**

4. Next, transfer 10 µl from the second well to the third well, mix with a pipettor. This is a $10^{-2}$ dilution.

5. Continue with the dilutions until you have finished with a $10^{-4}$ dilution (the 5th well, check image below).

6. Using the grid on the next page, spot **20 µl** of each well onto each of the 4 **LB** plates (0 IBU, 10 IBU, 50 IBU, 100 IBU).

7. Continue with *L. buchneri*, briefly vortex the culture tube, and transfer 100 µl to the first well of the **MRS** row. **Note: make sure that *L. buchneri* is added to MRS and not LB media.**

8. Again, continue with the dilutions until you have finished with a $10^{-4}$ dilution

9. Using the plate grid, spot **<u>4 µl</u>** of each well onto each of the 4 **MRS** plates (0 IBU, 10 IBU, 50 IBU, 100 IBU).

10. Incubate plates 1-3 days at 37°C, take a picture of the plates, and qualitatively score and record sensitivity (e.g., no sensitivity equals '+', partial sensitivity equals '+/-', complete sensitivity equals '-').

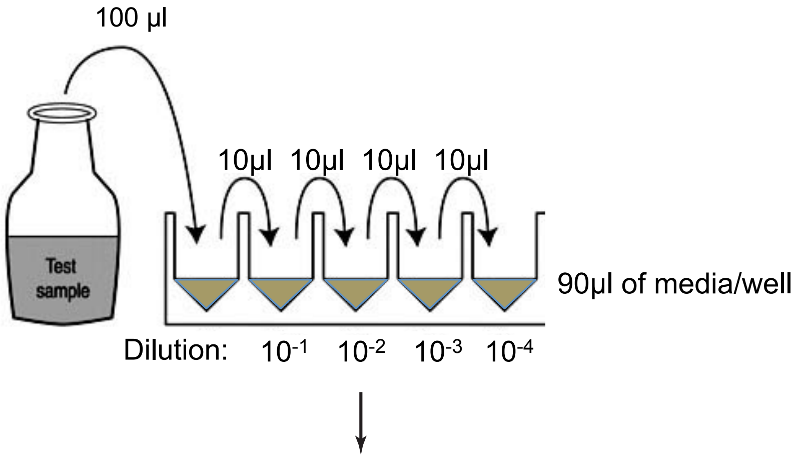


Spot 20 µl (*E. coli*) or 4 µl (*L. buchneri*) of each well onto the control and hop extract plates. Use the grid below as a guide: place plate on top of grid, and pipet each well into a square (i.e., well #1 pipetted into square #1, well #2 pipetted into square #2, etc.)

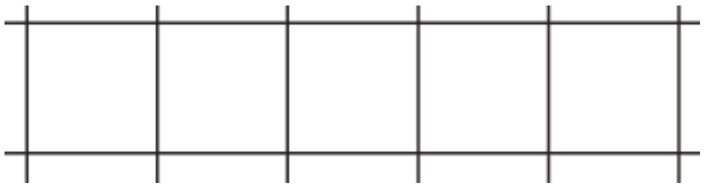

**<u>Recipes:</u>**

<u>Lysogeny Broth (LB) per liter</u>
yeast extract........................5 g
tryptone...............................10 g
NaCl...................................5 g

<u>de Man, Rogosa, and Sharpe (MRS) Broth per liter</u>
proteose peptone No. 3........5 g
beef extract..........................10 g
yeast extract.........................5 g
dextrose...............................20 g
polysorbate 80......................1 g
ammonium citrate.................2 g
sodium acetate......................5 g
magnesium sulfate................0.1 g
manganese sulfate.................0.05 g
dipotassium phosphate..........2 g
